# Effects of age on resting-state cortical networks

**DOI:** 10.1101/2024.09.23.614004

**Authors:** Chetan Gohil, Oliver Kohl, Jemma Pitt, Mats W.J. van Es, Andrew J. Quinn, Diego Vidaurre, Martin R. Turner, Anna C. Nobre, Mark W. Woolrich

## Abstract

Understanding how ageing affects brain function remains a central challenge in neuroscience. Electrophysiological brain imaging techniques provide a near-direct measure of neuronal activity, which is useful for characterising neurophysiological health. They offer us the ability to track large-scale networks of functional activity with high temporal precision. The effects of healthy ageing on these networks remain poorly understood, in part due to small sample sizes and limited control for confounding factors in previous studies. Here, we analysed resting-state source-reconstructed magnetoencephalography (MEG) data from a large cross-sectional cohort of healthy adults (*N* =612, 18-88 years old) to characterise the effect of age using not only time-averaged (static), but also transient (dynamic) network activity. We examined time-averaged power and coherence across canonical frequency bands (*δ, θ, α, β, γ*), as well as transient network dynamics identified using Hidden Markov Modelling. We included many confounding variables known to be affected by age, such as brain volume, as well as head size and position, which have previously been overlooked. Ageing was associated with frequency-specific changes in oscillatory power, with decreases in low-frequency (*δ, θ*) power and increases in high-frequency (*β*) power. Coherence increased across all frequency bands and was positively associated with cognitive performance. Transient network analyses additionally revealed that frontal network occurrences declined with age, with evidence suggesting a compensatory role in supporting cognition. These findings provide a more comprehensive electrophysiological signature for healthy ageing and establish a baseline for detecting pathological change.

**Highlights:**

- Using source reconstructed magnetoencephalography data, we studied cortical functional networks of oscillatory activity from a large healthy population (612 subjects, 18-88 years old).
- Time-averaged (static) network analysis in canonical frequency bands (*δ, θ, α, β, γ*) showed: *β* power and coherence increased with age; δ/*θ* power decreased with age, but coherence increased. Coherence in all frequencies increased with cognitive performance.
- Transient (dynamic) network analysis showed frontal networks decreased in occurrence with age and their relation to cognitive performance showed that these changes were compensatory. The other networks showed a decreased occurrence with age.
- In testing age and cognitive performance effects for statistical significance, we included a comprehensive set of confounds, such as brain volumes, head size and position, which has previously been overlooked.
- We provide a comprehensive electrophysiological signature for healthy ageing and establish a baseline for detecting pathological change.

## 1 Introduction

With an increasing proportion of elderly people across the globe [1], there is a pressing need to understand age-related changes in the brain. It is important to separate changes associated with healthy cognitive decline from those linked to abnormal cognitive decline or specific age-associated diseases. This could facilitate the early detection of pathology and the development of preventative intervention [2, 3].

Ageing causes several multifaceted changes to the brain [4]. In this work, we focus on one aspect of these changes, namely the effect^1^ of age on functional brain activity. Previous work has predominantly characterised this using functional magnetic resonance imaging (fMRI) [5]. However, electrophysiological recordings of brain activity, such as magneto/electroencephalography (M/EEG), offer a unique perspective on brain function by providing us a direct measure of neuronal activity at its natural timescale [6].

An observed property of the functional activity from neuronal populations and circuits is the emergence of oscillations [8]. Our understanding of brain function is rapidly moving towards a whole-brain description [9], where cognition is facilitated by large-scale functional networks [10, 11]. Neuronal oscillations are believed to help regulate the routing of information in these networks [12, 13]. Note that functional network activity is inherently transient in nature [14]. MEG can be used to reliably estimate large-scale cortical networks of transient oscillatory activity at millisecond timescales [14, 15, 16].

Neuronal oscillations reflect the underlying neurophysiology of the brain [6]. Consequently, pathological changes in the neurophysiology, e.g. the synaptic health, may be detectable via alterations to large-scale networks of oscillatory activity. To detect such changes, we must first characterise a healthy state as a function of age. In this work, we make progress towards this by studying the effect of age on cortical networks of oscillatory activity.

In this work, we will estimate *source* activity within the cortex using resting-state MEG recordings. We focus on resting-state MEG because it captures brain activity without relying on a specific task or performance, which can vary with age. Resting-state paradigms are also well-suited for elderly and clinical populations and can be collected at scale. Recent studies that have looked at age effects in source localised MEG have focused on *time-averaged* networks [17], and few have looked at network dynamics. Here, we will study a more comprehensive description of age effects on functional networks. We will look at both time-averaged and transient networks using one of the largest cohorts to date (612 subjects, cross sectional, 18-88 years old). We will perform an exploratory analysis to identify the network features that can be linked to age and cognition in Section 3 and discuss our findings in the context of previous neuroimaging studies in Section 4. We will also include a comprehensive set of confounding variables in testing for statistical significance, such as brain volumes, head size and position, which have been overlooked in previous studies.

## 2 Materials and Methods

### 2.1 Dataset

We study a cross-sectional group of 612^2^ healthy subjects aged between 18 and 88 years from the Cam-CAN (Cambridge Centre for Ageing and Neuroscience) dataset. The demographics of these subjects are shown in Figure 1. Each subject has a resting-state^3^ (eyes closed) MEG recording, structural MRI (sMRI) and a set of cognitive task scores. Participants were instructed to remain awake throughout the ∼8-minute resting-state acquisition, although no independent physiological measures of vigilance were obtained. Further information regarding the dataset and protocols is provided in [18, 19].

**Figure 1:**
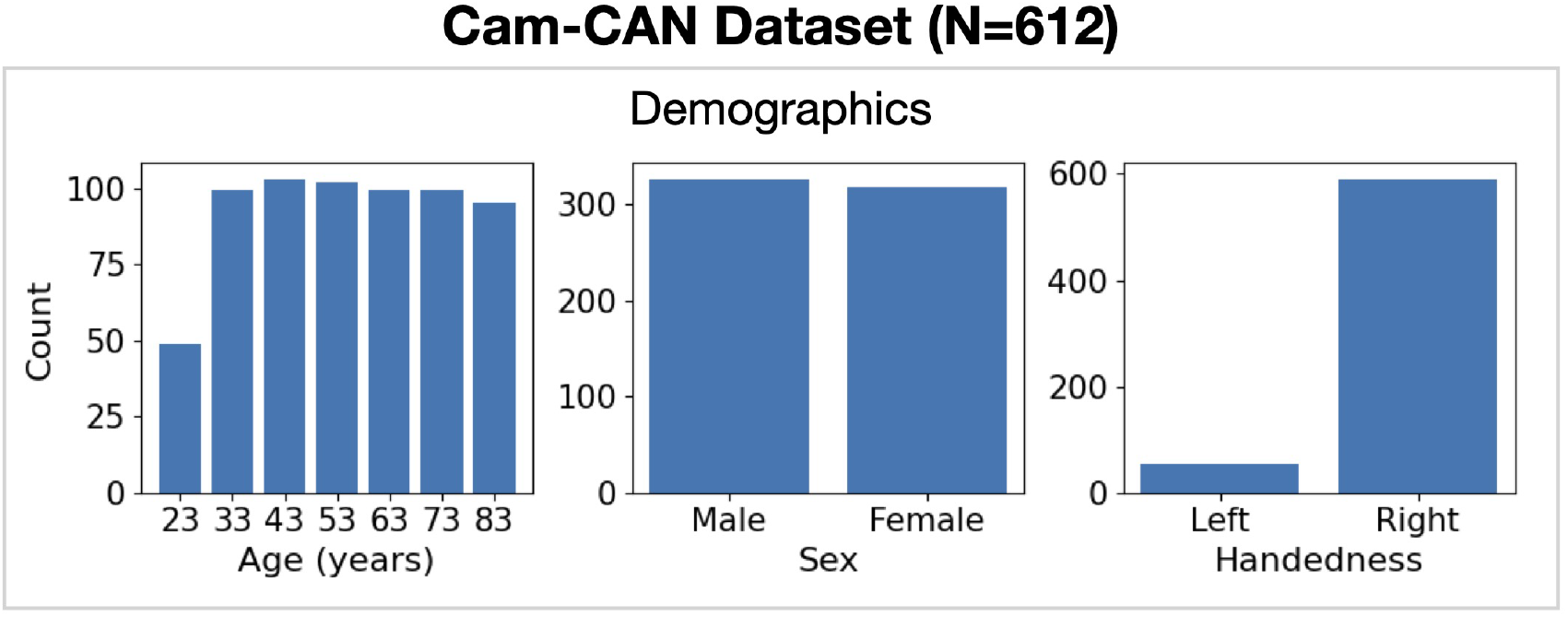
Demographics. Distribution of ages (left); sex (middle) and handedness (right).

#### MEG data

The MEG recordings were obtained using a 306 channel Vectorview system (Elekta Neuromag, Helsinki, Finland), consisting of 102 magnetometers and 204 orthogonal planar gradiometers. The acquisition time of the recordings was 8 minutes 40 seconds, with the first 20 seconds discarded. The data were recorded at a sampling frequency of 1 kHz and bandpass filtered between 0.03 Hz and 330 Hz.

#### sMRI data

T1-weighted sMRIs were acquired using a 4 minute and 32 second MPRAGE (Magnetization-Prepared RApid Gradient Echos) sequence with a 3 T TIM Trio scanner (Siemens Healthcare, Erlangen, Germany) equipped with a 32-channel head coil.

#### Cognitive task scores

Five broad cognitive domains were evaluated using a set of tasks (executive function, language, emotion, memory, motor control). We drew from the same 13 cognitive task scores as [56], provided by [21].

We reduced the cognitive task scores into a single measure for performance by concatenating the scores into a vector for each subject and applying Principal Component Analysis (PCA) across subjects. We used the 10 cognitive tasks summarised in Table 1. Three cognitive tasks (Face Recognition, Spot the Word, Proverb Comprehension) were excluded due to their incompatibility with PCA for dimensionality reduction (their distribution was highly non-Gaussian or discrete). Taking the first principal component gives a single number that quantifies the performance of a subject across all of the cognitive tasks. Figure 2 shows the PCA loadings (with the contribution of each cognitive task to the principal component). All PCA loadings have positive values, indicating an increase in the first principal component results in an increase in all cognitive task scores.

**Table 1:**
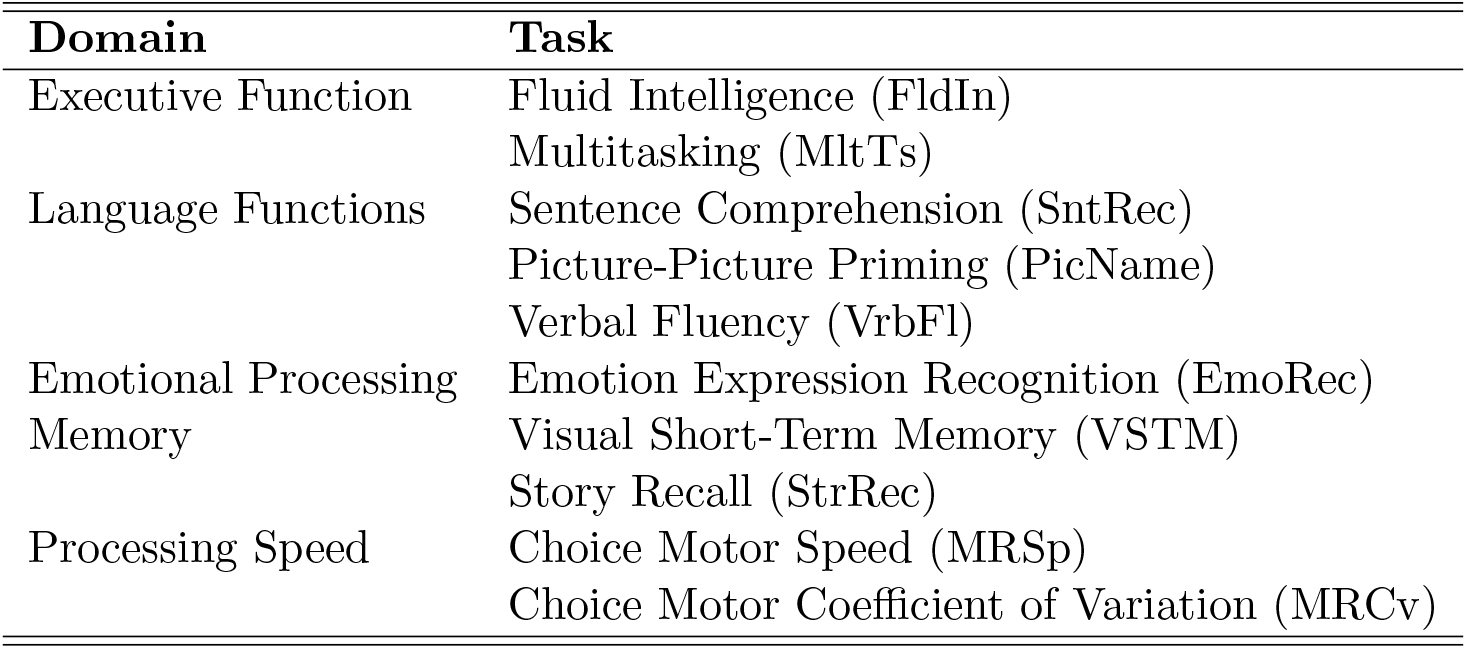
Tasks used for evaluating cognitive performance. See Borgeest et al. (2018) for a detailed description of each task.

**Figure 2:**
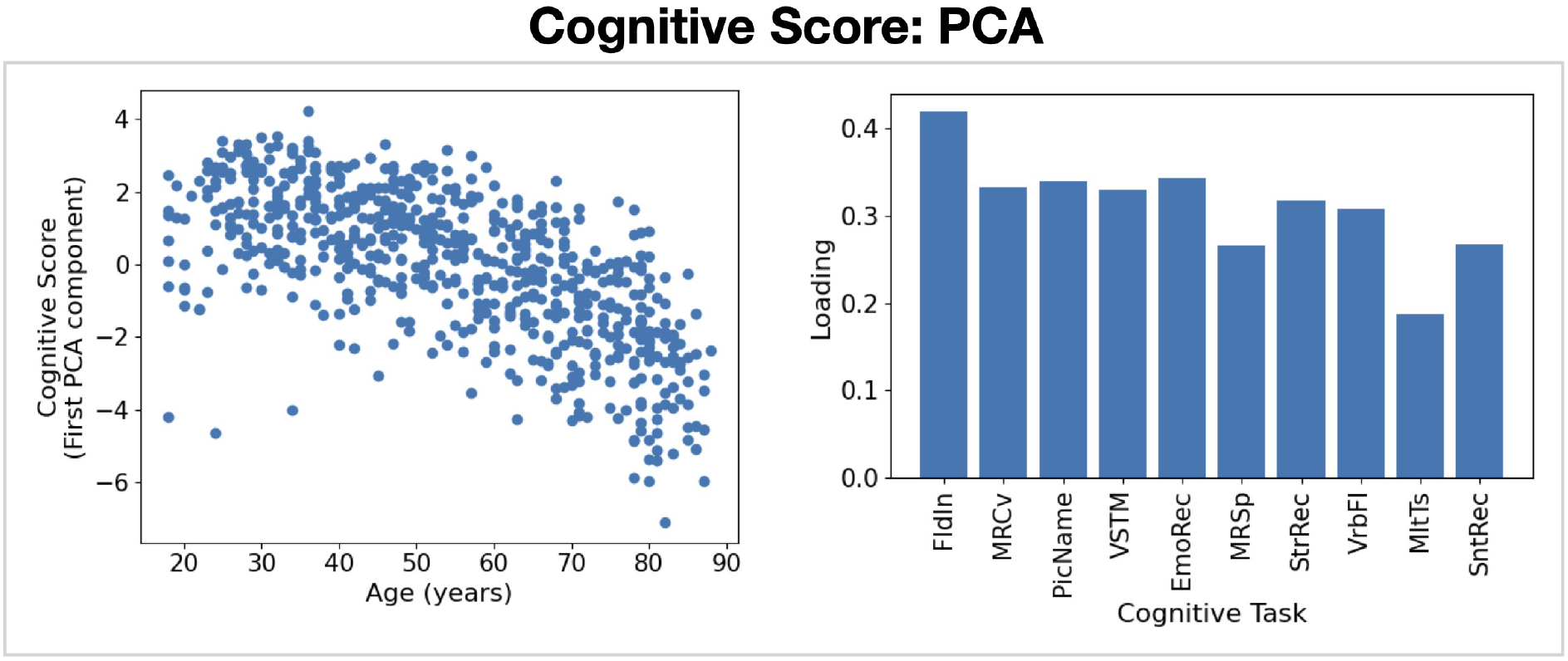
PCA applied to the cognition scores. Correlation of the first principal component with age (left) and PCA loadings (right). See Table 1 for the definition of the acronym for each task.

#### 2.1.1 MEG Preprocessing, Source Reconstruction and Parcellation

The Cam-CAN MEG data were processed using the `osl-ephys` toolbox [22, 23, 24, 25]. A detailed description of the preprocessing, source reconstruction and parcellation is provided in SI Section 1.1. In brief, to preprocess the sensor-level data for each subject we applied a tSSS MaxFilter [26], downsampled to 250 Hz, band-pass filtered to 1-80 Hz, applied automated bad channel and segment detection, automated independent component analysis (ICA) cleaning to remove ocular/cardiac noise and interpolated bad channels. Following this, we extracted the brain and skull surface from a sMRI and coregistered the MEG [23]. Source reconstruction was performed using a volumetric LCMV beamformer and the voxel-level data was parcellated using a 52 region-of-interest (ROI) atlas [27] using principal component analysis (PCA) applied to the voxel-level data. Finally, the parcel time courses were orthogonalised [28], ‘sign flipped’ [14], and standardised (z-scored) temporally. Note, in the current work all references to the ‘parcel time courses’ correspond to the standardised sign-flipped parcel time courses.

### 2.2 Time-Averaged Network Analysis

We assessed the impact of healthy biological ageing on the following time-averaged properties of source-localised MEG data: power spectral density (PSD), power, and coherence. The calculation of these quantities is described below. Figure 3 summarises the steps. Note, although we band-pass filtered the data between 1-80 Hz, we will focus on the range 1-45 Hz because this is the relevant frequency range for large-scale networks [29].

**Figure 3:**
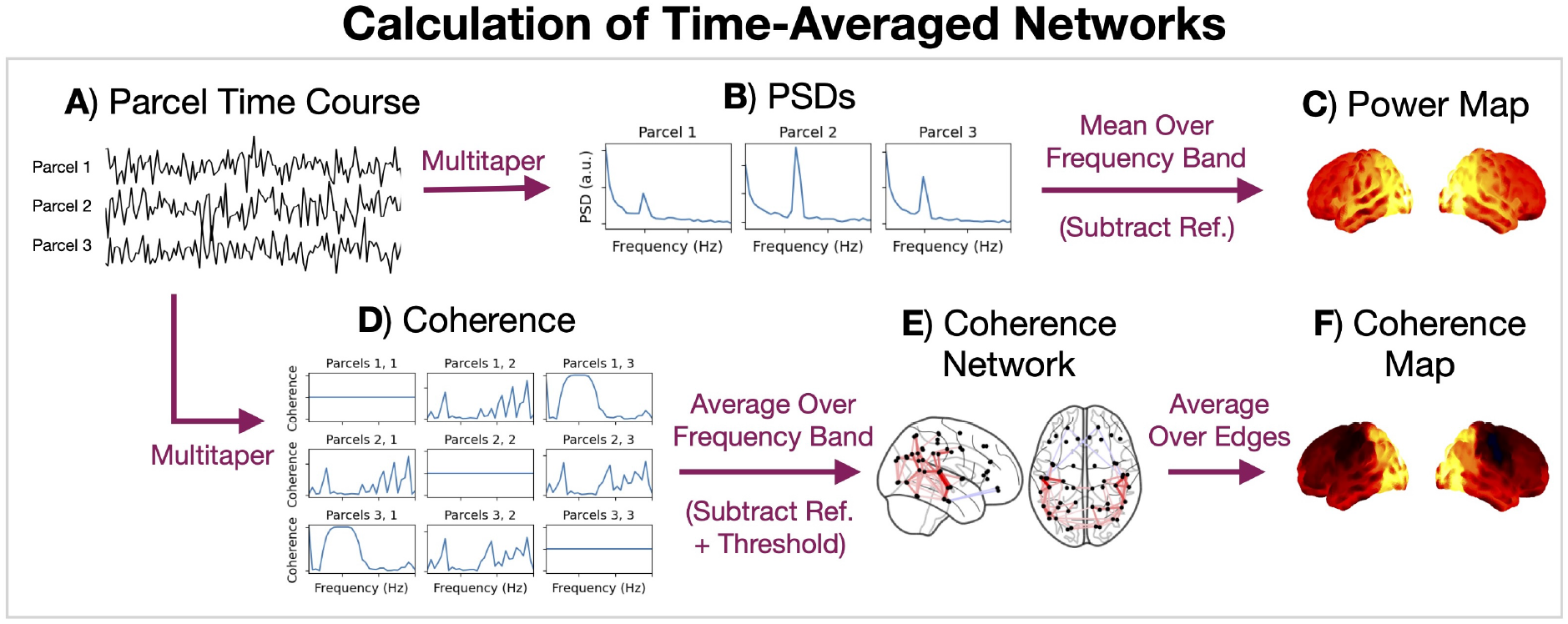
Calculation of time-averaged networks for an individual subject. Standardised parcel time courses (A) are used to calculate power spectral densities (PSDs, B) for each parcel using the multitaper method. The PSDs are averaged over a frequency band to give the power at each parcel, which is visualised as a brain surface heat map (C). The parcel time courses are also used to calculate cross spectral densities, which are normalised to give the coherence (D) for each pairwise combination of parcels. These are averaged over a frequency band and visualised as a graphical network (E). Averaging over the (unthresholded) network edges for a given parcel, we get a brain surface heat map (F).

#### Power/coherence spectra

We used the multitaper method (2 s window, 0% overlap, 7 DPSS tapers, 4 Hz time half bandwidth) to calculate a PSD for each parcel and a cross spectral density (CSD) for each pairwise combination of parcels. The multitaper parameters used in this work are typical in MEG analysis [40]. We did this for each subject individually and (temporally) standardised (z-scored) the parcel time courses before calculating the P/CSDs^4^. We obtained a (subjects, parcels, frequencies) array for the PSDs and (subjects, parcels, parcels, frequencies) array for the CSDs.

#### Canonical frequency bands

The P/CSD is a function of frequency. In time-averaged MEG analysis, it is common to reduce the frequency dimension by studying the activity in a particular frequency band. In the current work, we look at the following frequency bands: *δ* (1-4 Hz), *θ* (4-8 Hz), *α* (8-13 Hz), *β* (13-24 Hz), *γ* (30-45 Hz). We have chosen the frequency range 13-24 Hz for *β* due to a site-specific artefact observed in the coherence in the range 24-30 Hz that could not be removed with notch filters.

#### Power maps

Averaging a PSD over a frequency band, we get a *power density*, which we simply refer to as the *power* in the frequency band. We calculated the power at each parcel for each subject by averaging a PSD within the five canonical frequency bands described above. This resulted in a [subjects, parcels] array for each frequency band.

Each [parcels,1] array is referred to as a *power map*, which we can visualise as a heat map plot on the brain surface. Normally, when visualising a power map, we are interested in looking at the power relative to a chosen reference. In the time-averaged power analysis, we used the weighted (parcel-specific) average across frequency bands as the reference. This shows the spatial pattern of band-specific power relative to the power across all frequency bands. This was only done in the visualisation of group-averaged power maps.

#### Coherence networks/maps

We calculated the coherence using the P/CSD for each pairwise combination of parcels for each subject as

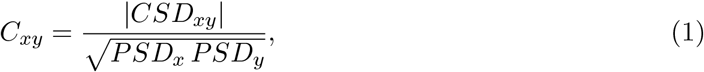

where |.| denotes the absolute value and *x, y* indicate different parcels. This resulted in a [subjects, parcels, parcels, frequencies] array. Averaging over the frequency dimension, we obtained a [subjects, parcels, parcels] array for each canonical frequency band.

Each [parcels, parcels] array represents a graphical network. The off-diagonal elements in the (parcels, parcels) array represent *edges* in the network. The graphical network was visualised by first subtracting a reference, for which we used the (edge-specific) weighted average across frequency bands, then thresholding to show the top 3% irrespective of sign. The colour of each edge indicates its value. The centroid of each parcel was used for the location of each node in the network.

We calculated a *coherence map* by averaging over the edges for each parcel, which resulted in a [parcels,1] array that can be visualised in the same way as a power map. We used the original [parcels, parcels] array before subtracting the reference and thresholding for this.

### 2.3 Transient Network Analysis

We used the method introduced in [14] to learn transient networks of coherent activity in source localised MEG data. This method applies a machine learning approach for segmenting time series data, known as the Hidden Markov Model (HMM, described below), on time-delay embedded (TDE)-PCA data. Applying the HMM to TDE data is an approach that allows us to model dynamics in the oscillatory activity in the data [14, 15]. Figure 4 summarises the calculation of transient networks using this approach. We describe each step below.

**Figure 4:**
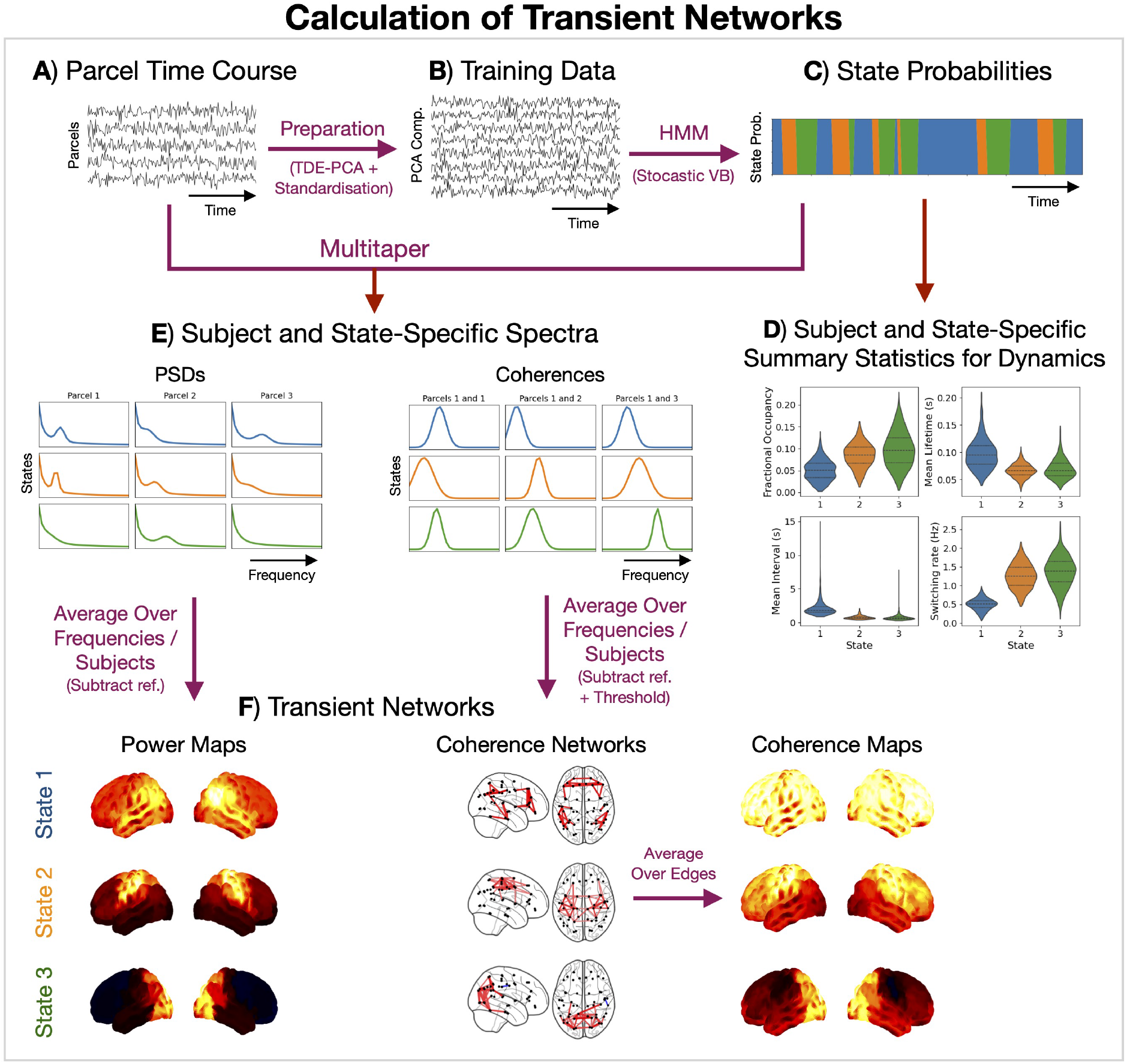
Calculation of transient networks using a group-level HMM. Parcel time courses (A) are prepared by performing time-delay embeddings (TDE), principal component analysis (PCA) and temporal standardisation (z-scored) to give the data we train the HMM on (B). Using a stochastic variational Bayes algorithm, we infer the probability of each hidden state at each time point in the training data (C). We summarise the dynamics of each subject and state by calculating statistics from the state probabilities (D). The HMM is trained at the group level. Using the parcel time course and inferred state probabilities we *dual estimate* the power spectral density (PSD) and coherence for each subject and state (E). Averaging these over a frequency band, we get a power map and coherence network/map for each state (F). For visualisation we show the power maps and coherence networks relative to the mean across states and threshold the coherence networks to show the top 3% irrespective of sign. The coherence maps are calculated using the unthresholded coherence networks and shown relative to the mean across states. Only 3 states have been shown in this figure for illustration.

#### 2.3.1 Data Preparation

Before applying the HMM, the parcel time courses were prepared by performing TDE, which augments the parcel data with extra channels containing time-lagged versions of the original parcel data. In the current work, we used ±7 lags (±28 ms window), resulting in 780 channels. PCA was then applied at the group level to reduce the TDE data down to 120 channels. The selection of the TDE and PCA parameters is discussed in [15]. Finally, we (temporally) standardised (z-scored) the TDE-PCA data. These were the data used to train the HMM.

#### 2.3.2 The Hidden Markov Model

The HMM [30, 31] is a generative model for time series data. At each time point, *t*, there is an underlying categorical hidden state, *θ*_*t*_ ∈ {1, …, *K*}, where *K* is the number of states. The observed time series, ***x***_*t*_, is generated using a multivariate Normal distribution based on the hidden state:

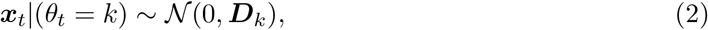

where *θ*_*t*_ = *k* is the hidden state at time point *t* and ***D***_*k*_ is a *state covariance*. Note, we force the mean to be zero to focus on modelling dynamics in the covariance, i.e. the connectivity information we are interested in. Dynamics are governed by transitions in the hidden state. Note, it is assumed that the probability of a transition at time point *t* only depends on the state at the previous time point *θ*_*t*−1_ (this is the Markovian constraint). Each pairwise state transition probability is contained in the *transition probability matrix*:

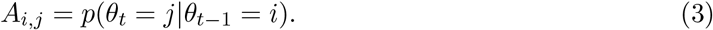

When we train an HMM on data, we learn the most likely value for the state covariances, {***D***_1_, …, ***D***_*K*_}, and transition probability matrix, ***A***, to have generated the data. We use stochastic variational Bayes [31] to do this, where we iteratively update our estimates for {***D***_1_, …, ***D***_*K*_, ***A***} based on a random subset of the training data to minimise the variational free energy. We use the Baum-Welch algorithm [31] to calculate the (*posterior*) probability of each state being active at each time point, *q*(*θ*_*t*_), for each subject based on our estimates for {***D***_1_, …, ***D***_*K*_, ***A***}. The posterior, *q*(*θ*_*t*_), is a (states, time) array for each subject.

#### Hyperparameters

We used the HMM implemented in the `osl-dynamics` toolbox [15]. To train an HMM, we need to specify a few hyperparameters. An important hyperparameters is the number of states. In the current work, we identified 10 states for comparability with previous HMM MEG studies [14, 15, 32, 33, 34, 35, 36, 37, 38, 39, 40, 41], which used 8-12 states. Other hyperparameters include the batch size, sequence length and learning rates. These had little impact on the HMM inference. The values used are summarised in Table S1.

#### Run-to-run variability

The final estimates for {***D***_1_, …, ***D***_*K*_, ***A***} can be sensitive to the initial values used at the start of training. The typical approach for overcoming this is to train several models from scratch starting from random initialisation and picking the one with the lowest final variational free energy (deemed the best description of the data) for the subsequent analysis. Historically, this has produced very reproducible results [15]. In our case, we analyse a particularly large dataset. This makes the HMM inference very stable, and we consistently converge on very similar estimates for {***D***_1_, …, ***D***_*K*_, ***A***}. Despite this, we took a cautionary approach and selected the best model from a set of five runs for analysis.

#### 2.3.3 Post-Hoc Analysis

Once we trained an HMM and obtained a state probability time course for each subject, *q*(*θ*_*t*_), we calculated a *state time course* by taking the most probable state (*maximum a posteriori probability estimate*) at each time point, 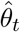. We then performed post-hoc analyses for each subject using 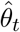. Note, 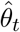 is mutually exclusive.

##### Summary statistics for state dynamics

We summarise the dynamics of each transient network by calculating summary statistics based on its state time course. For each subject and state, we calculated:

- Fractional occupancy: the fraction of total time spent in a state.
- Mean lifetime (ms): the average duration a state was active.
- Mean interval (s): the average duration between successive state activations.
- Switching rate (Hz): the average number of state activations per second.

These summary statistics are highly correlated. We will sometimes refer to them all jointly as the *occurrence*. An ‘increased occurrence’ refers to an increase in fractional occupancy, mean lifetime, switching rate and decrease in mean interval jointly.

##### Transition probabilities

We calculated a subject-specific transition probability matrix by counting the number of pairwise transitions for each combination of states in the state time course and normalising to ensure a sum-to-one constraint.

##### State power/coherence spectra

Each HMM state represents a transient network of characterised by distinct spectral (e.g. power spectra and coherence) activity. Here, we combined the state time course with the original parcel data (pre-TDE-PCA) to estimate the spectral properties (PSD and coherence) of each state using a multitaper [42]. This involved performing the following steps for each subject and state:

1. The (temporally) standardised parcel time courses are multiplied by the state time course.
2. The P/CSD for each parcel/pair of parcels us calculated using a multitaper (2 s window, 0% overlap, 7 DPSS tapers, 4 Hz time half bandwidth). This is the standard approach/settings used for HMMs trained on TDE-PCA data [14, 40].
3. Equation (1) is used to calculate the coherence from the P/CSD.
4. The amplitude of the PSD is biased by the amount of time each state is active due to the multiplication in step 1. We accounted for this by scaling the PSD by one over the fractional occupancy of the state.

##### State power maps

The multitaper results in a [subjects, states, parcels, frequencies] array containing the PSDs. Similar to the time-averaged power analysis, we need to reduce the frequency dimension by averaging over a band. The HMM’s objective is to identify spectrally distinct states, i.e. temporally segment occurrences of different oscillatory activity. Therefore, each HMM state tends to have its own characteristic PSD. This means we do not need to specify the frequency band by hand and can integrate over the full frequency range the of the PSD. Note, in practice we only calculated the PSD for the frequency range 1-80 Hz due to the bandpass filter we applied before source localisation. Averaging over the frequency dimension results in a [subjects, states, parcels] array. In the visualisation of the group-averaged power maps, we displayed each state’s power map separately and used the average (parcel-specific) power across states as the reference.

##### State coherence networks/maps

The multitaper results in a [subjects, states, parcels, parcels, frequencies] array for the coherences. Similar to the state power maps, we reduce the frequency dimension by taking the average across the full frequency range (1-80 Hz). This results in a [subjects, states, parcels, parcels] array, which represent the state coherence networks. These are visualised in the same way as the time-averaged coherence networks. We displayed the group-average coherence network for each state individually using the (edge-specific) average across states as the reference and thresholding the top 3% of edges irrespective of sign. The state coherence maps were calculated in the same way as the time-averaged coherence map using the non-referenced, unthresholded state coherence networks.

### 2.4 Statistical Significance Testing

In the current work, we characterised the impact of healthy ageing on resting-state functional networks by using a General Linear Model (GLM) with an age regressor to predict summary target features calculated from the networks. The summary target features used were:

- The power and coherence within canonical frequency bands from time-averaged networks (described in Section 2.2);
- The power, coherence and summary statistics for dynamics for the transient networks identified by the HMM (described in Section 2.3)

#### GLM permutations

We employed non-parametric permutations with a GLM to test for statistical significance. This approach is described in detail in [43]. It involves fitting a group-level model:

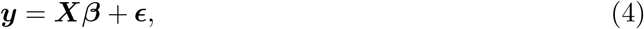

where ***y*** [subjects, features] is the target data, ***X*** [subjects, regressors] are regressors, referred to as the *design matrix*, ***β*** [regressors, features] are regression coefficients, referred to as *parameter estimates* or *effects*, and ***E*** [subjects, features] are the residuals.

For a given ***X***, fitting the group-level GLM to the data ***y*** provides an (ordinary-least-squares) estimate of ***β*** and ***E***. The effect ***β***_*i*_ for regressor ***X***_*i*_ indicates how ***y*** would change with ***X***_*i*_. ***β***_*i*_ is a [features,1] array and ***X***_*i*_ is a [subjects,1] array.

Statistical significance testing involves selecting a regression coefficient of interest and building a null distribution of test statistics based on that coefficient. We do this by repeatedly permuting the design matrix and refitting the GLM. We describe this in more detail below.

#### 2.4.1 Age Effects

We are particularly interested in correlations with age, i.e. ***β***_*i*_ for when *i* = Age. We refer to ***β***_Age_ as the *age effect*.

##### Confounds

Isolating the change in functional brain activity due to age is difficult due to the many confounding variables that change with age and the heterogeneity in functional brain activity across subjects. Here, we include a comprehensive set of confounds: sex, total brain volume, relative grey matter volume, relative white matter volume, head size, (*x, y, z*) head position, and the PCA-reduced cognitive performance score. The grey/white volumes are relative to the total brain volume. The brain volumes were calculated from the sMRIs using FSL’s anatomical processing script (`fsl_anat`). The head size and position were estimated by fitting a sphere to the Polhemus head shape points and fiducials. The design matrix used to study age effects is shown in Figure S1. We standardise each regressor (z-scored across the subject dimension) and include a constant (mean) regressor.

##### Statistical significance testing

We tested whether the age regression coefficient differed significantly from zero:

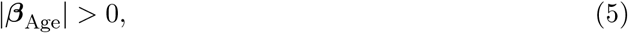

where |.| denotes the absolute value. We built a null distribution for ***β***_Age_ by randomly permuting the design matrix ***X***, refitting the GLM, and recording the maximum absolute regression coefficient across all features (i.e. max(|***β***_Age_|)) as the test statistic.

Importantly, for each permutation the GLM was fit once with all features included simultaneously in ***y***, and the same permuted regressor was applied to every feature. Thus, a single null distribution of maximum statistics was constructed across features, rather than separate null distributions for each feature. In the brain-wide analyses, these features corresponded to all regions (and associated frequencies/states) analysed simultaneously, such that the maximum-statistic procedure controlled the family-wise error rate across the entire brain.

We used the absolute regression coefficient as the test statistic rather than a (one sample) *t*-statistic because the variance of each feature was very different. The high variance in particular features suppresses the *t*-statistic and sensitivity to effects in the features [43]. In all permutation testing we used the regression coefficient as the test statistic apart from when looking at linear age effects in the summary statistics for dynamics, where we used the *t*-statistic due to the different effect size in each feature^5^.

We applied 1,000 random ‘sign-flip’ permutations to the age regressor, where each entry in ***X***_Age_ had a 50% chance of being multiplied by -1. This resulted in a null distribution of size 1000. Looking up the percentile at which the observed absolute regression coefficient (calculated with the non-permuted design matrix) occurs in the null distribution gave us our *p*-value. We obtained a *p*-value for each feature. Sufficiently small *p*-values (*<* 0.05) were deemed to be significant.

#### 2.4.2 Cognitive Performance Effects

We were also interested in functional brain activity that correlates with cognitive performance. This is given by ***β***_Cog. Perf._, which is referred to as the *cognitive performance effect*.

##### Confounds

The PCA-reduced cognitive score (see Section 2.1) is negatively correlated with age (see Figure 2). We need to be sure that any effect we observed from the cognitive score is not simply indirectly due to the age effect. To do this, we included age as a confound regressor in the design matrix used to study the cognitive performance effect, see Figure S1. We also included the same confounds we did when we studied the linear age effect (brain volume and head size/position). Including the age regressor in the design matrix can be seen as a conservative approach as this is equivalent to regressing out the age effect from the target data (and other regressors).

##### Statistical significance testing

We tested whether the cognitive performance regression coefficient differed significantly from zero:

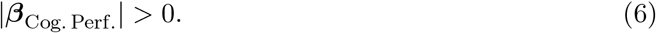

We used the same procedure for building the null distribution (1,000 sign-flip permutations) and calculating a *p*-value as we did for the linear age effect.

As above, each permutation involved fitting a single GLM across all features simultaneously and retaining the maximum absolute regression coefficient across features (brain wide: regions, frequencies, states) to form a family-wise error controlled null distribution. We used the regression coefficient as the test statistic in all cases apart from when predicting the summary statistics for dynamics, where we used the *t*-statistic instead.

## 3 Results

### 3.1 Time-Averaged Networks

We first characterised the time-averaged spatio-spectral properties of healthy individuals. Figure 5 shows the power spectral density (PSD), coherence spectrum, and time-averaged networks for different canonical frequency bands averaged over a large cohort of healthy individuals (*N* =612, 18-88 years old). We see healthy individuals exhibit unique spatial patterns of activity in each canonical frequency band.

**Figure 5:**
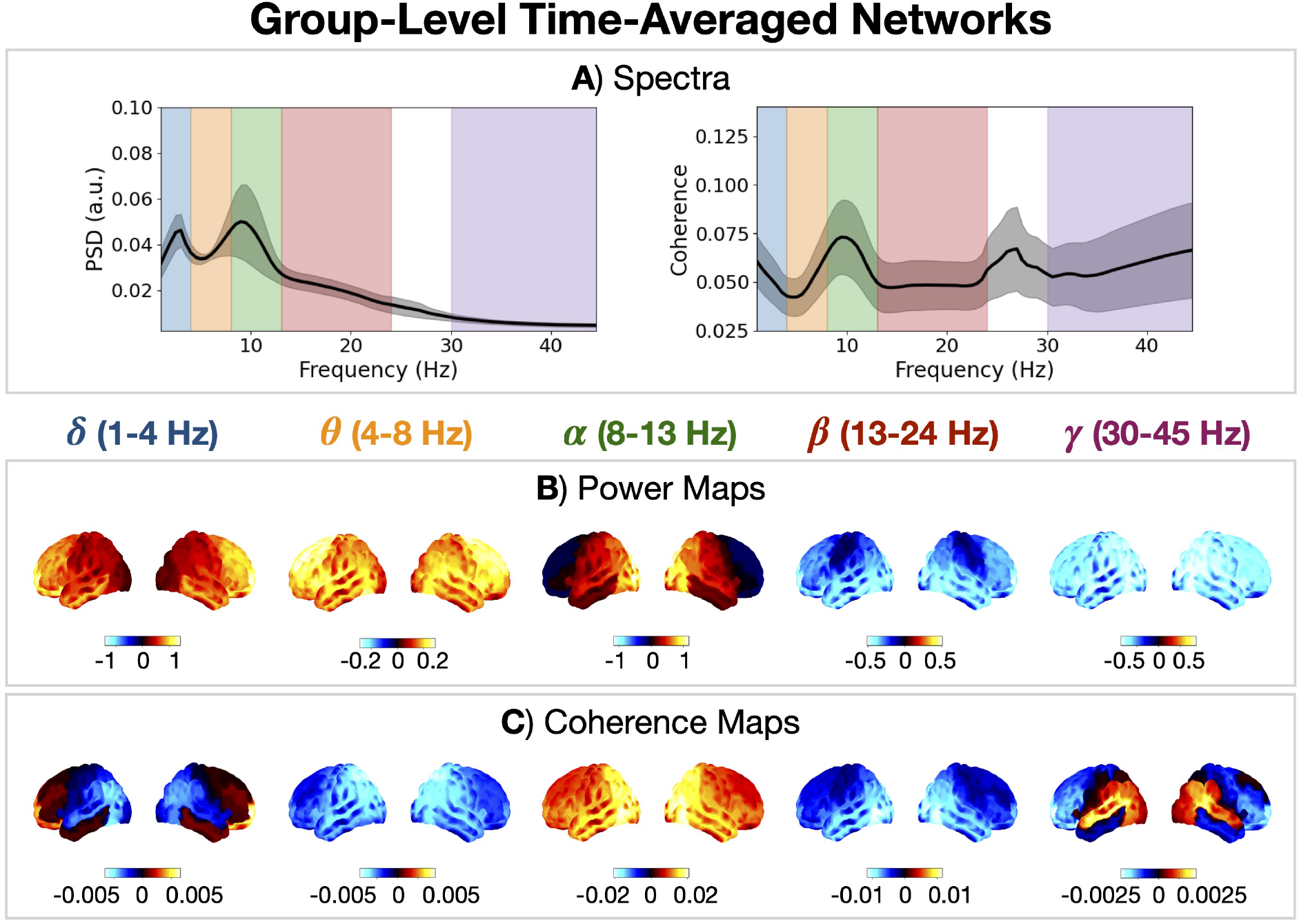
Healthy individuals exhibit frequency-specific networks of oscillatory activity. PSD (A, left) and coherence spectrum (A, right) averaged over parcels and subjects. The grey shaded area shows the standard deviation across subjects. Power (B) and coherence (C) maps for each canonical frequency band relative to the weighted mean across frequency bands. Unreferenced power maps are shown in Figure S2D.

The PSD decreases with frequency and there is a prominent peak in the *α* band, which indicates there are strong *α* oscillations present in the data (Figure 5A, left). Note, the dataset contained eyes closed resting-state MEG recordings, which are known to contain strong *α* activity [8]. In SI Section 1.2, we characterise the PSD with FOOOF (Fitting Oscillations and One Over F) [44]. The coherence spectrum is approximately flat except for the peak in the *α* band, which reflects phase locking between the *α* oscillations (Figure 5A, right). There is a site-specific artefact in the PSD and coherence spectrum between 24 and 30 Hz, which could not be removed with notch filters, therefore, this frequency range was not included in our definition of the *β*-band.

Figure 5B shows the spatial distribution of power in five canonical frequency bands (*δ, θ, α, β, γ*) and Figure 5C shows the coherence maps. The visualisation of the power and coherence maps highlights the activity unique to each band (the weighted mean across frequency bands was used as the reference). Low frequencies (*δ, θ*) have relatively high frontal power/coherence and low posterior coherence. The *α* band has relatively high occipital power and coherence. The *β* band has relatively high sensorimotor power and low posterior coherence. The *γ* band has low brain-wide power but high posterior coherence particularly in the temporal regions.

### 3.2 Age Effects in Time-Averaged Networks

Next, we characterised how the time-averaged network for each canonical frequency band was affected by age. Age effects in time-averaged power and coherence are shown in Figure 6. We see unique spectral changes occur with age for activity in each canonical frequency band.

**Figure 6:**
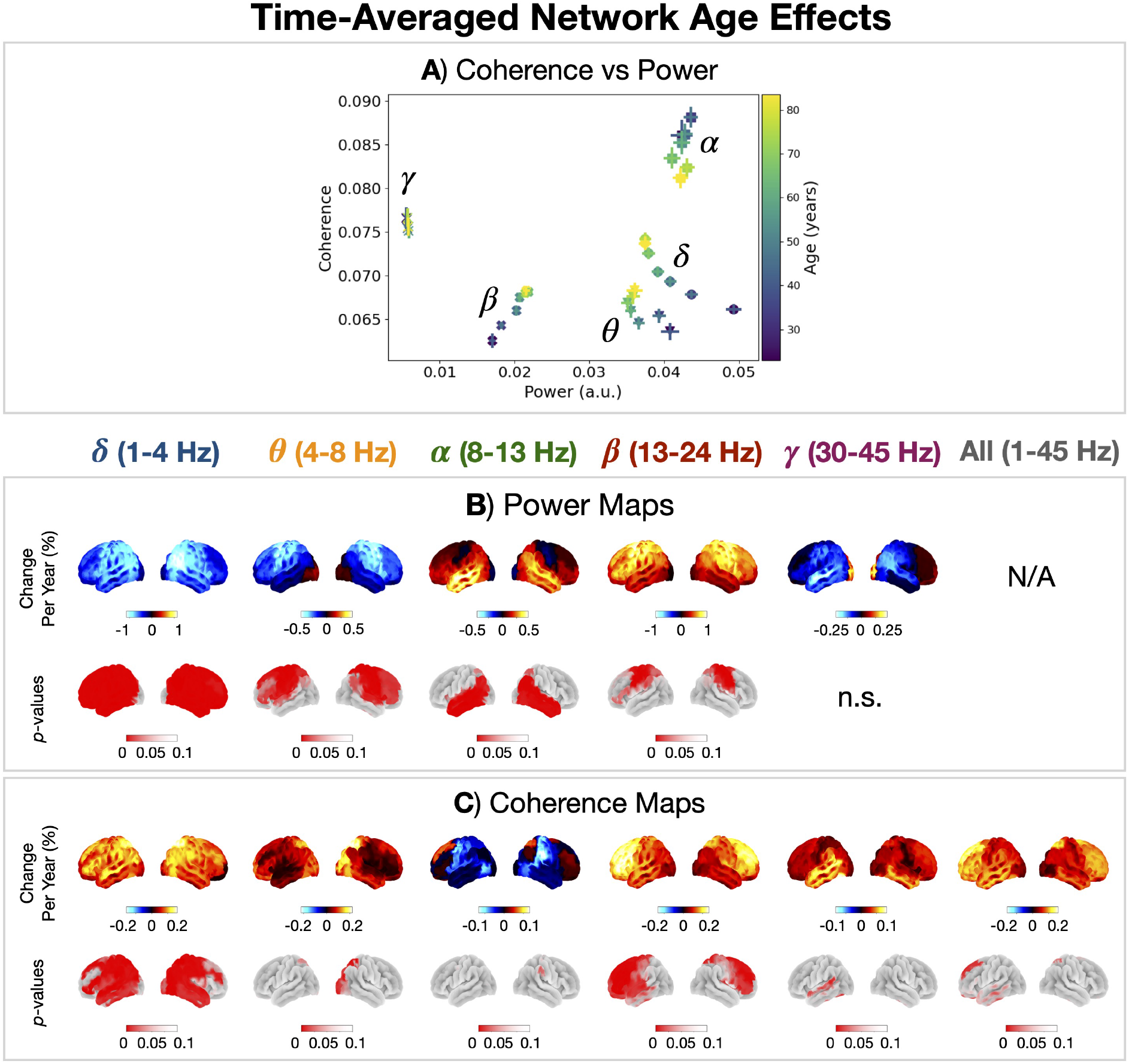
Unique spectral changes occur with age in the time-averaged networks for each canonical frequency band. A) Coherence vs power for each canonical frequency band averaged over frequencies, parcels and subjects for different 10 year cohorts (18-28, 28-38, …, 78-88 years old). The error bar shows the standard error on the mean. Age effects in power (B) and coherence (C) for each canonical frequency band.

The power and coherence in time-averaged networks follow a unique trajectory for each canonical frequency band (Figure 6A). Power in the *δ* and *θ* band decreases with age and coherence increases. Power in the *α* and *γ* remains stable with age but coherence decreases. Power and coherence in the *β* band increase with age.

Turning to the region-specific age effects in power (Figure 6B), there is: a decrease in brainwide *δ* power; increase in temporal-*α* power; increase in occipital-*θ* power, which reflects a slowing (leftward shift) of the *α* peak (Figure S2); and an increase in sensorimotor and frontal-*β* activity. Turning to the region-specific coherence age effects (Figure 6C), there is a general increase for all frequencies apart from the *α* band. Looking at the age effect in broadband (1-45 Hz) coherence, we see a greater increase in frontal (anterior) regions compared to posterior regions.

### 3.3 Cognitive Performance Effects in Time-Averaged Networks

Next, we identified the features of time-averaged networks that correlated with cognitive performance. Figure 7 shows cognitive performance effects in time-averaged networks. By accounting for age as a confound, we isolated the changes in power and coherence that could not be explained by age (or any other confound, such as brain volume or head size/position). Cognitive performance correlated with increased posterior-*α* power (Figure 7A) and a general increase in coherence for all frequencies (Figure 7B).

**Figure 7:**
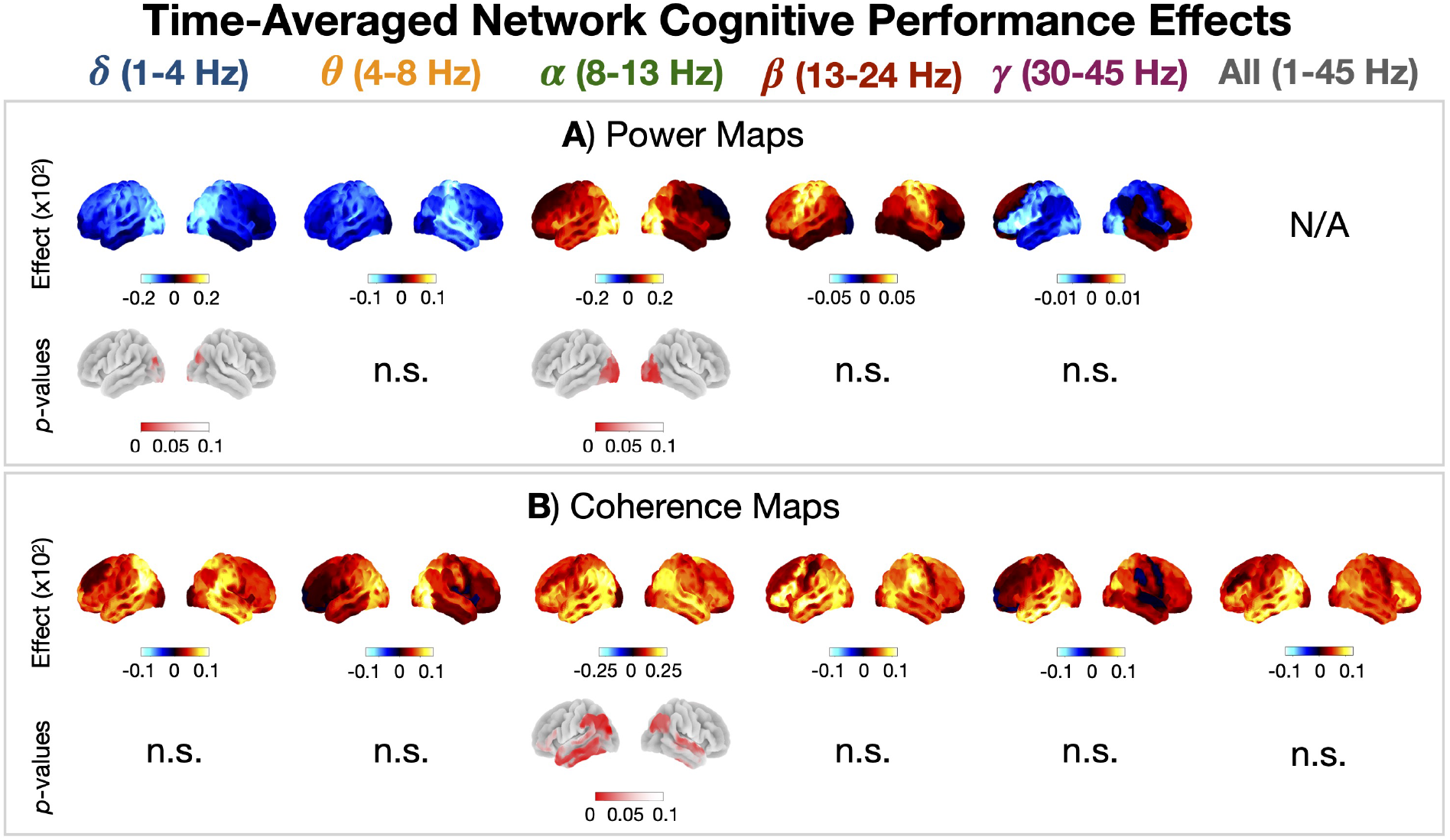
Time-averaged coherence correlates with cognitive performance. Cognitive performance effects in power (A) and coherence (B) for each canonical frequency band.

### 3.4 Transient Networks

Thus far, we have characterised brain activity using time-averaged (static) networks. However, time-averaged networks are a simplified, summary measure of what the brain is doing over time. In particular, brain networks have been shown to exhibit fast-switching dynamics [14, 15]. Figure 8 shows the result of using Hidden Markov Modelling to characterise these network dynamics in the form of ten transient network states that were inferred on the full cohort of healthy individuals (*N* =612, 18-88 years old). These transient networks have been found in multiple previous studies [14, 15]. However, this study has identified these networks in the largest cohort to date.

**Figure 8:**
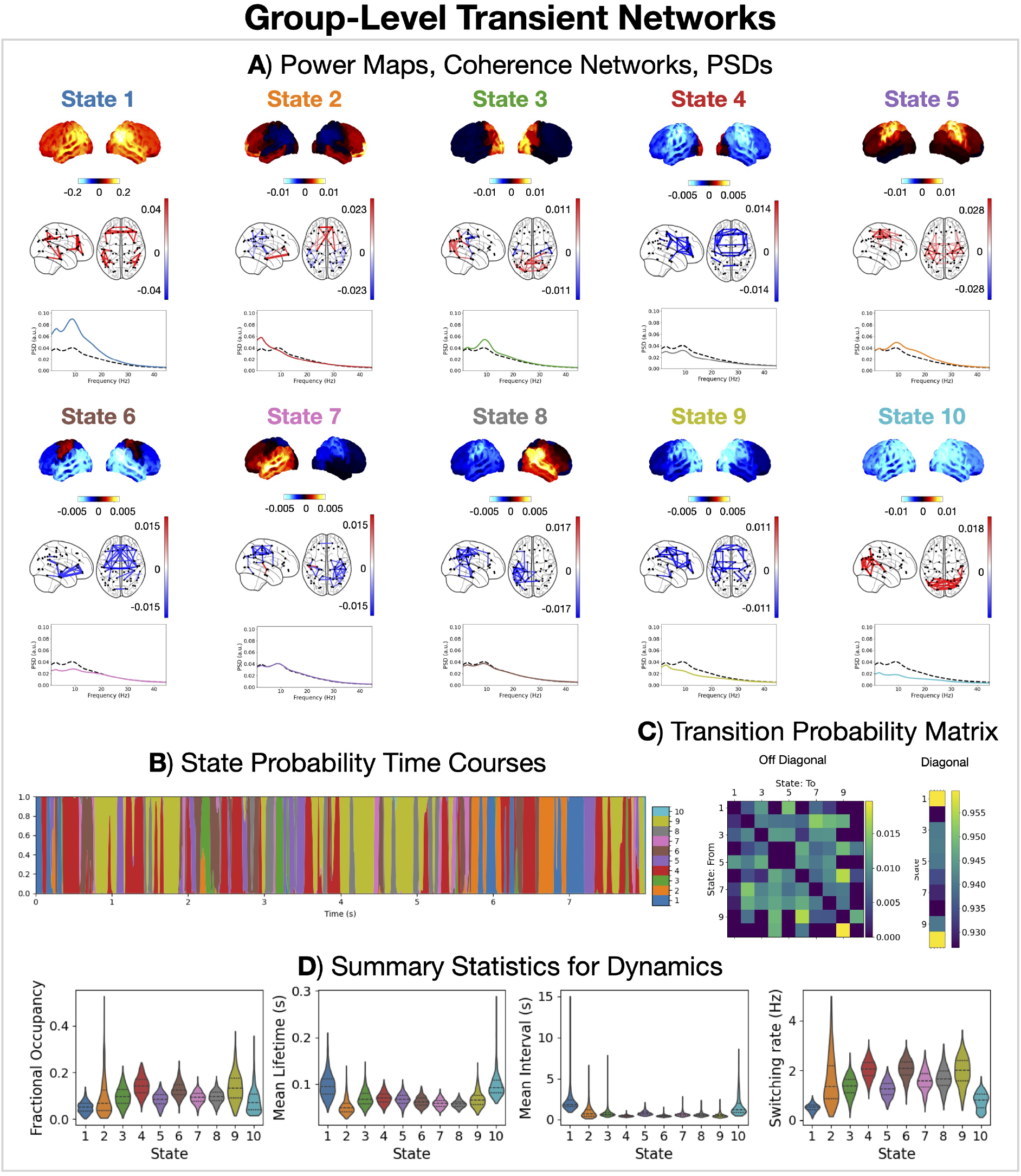
Healthy individuals exhibit fast (∼100 ms) transient networks of coherent activity. A) Power maps and coherence networks (top 3% of edges) averaged over subjects for each state and PSD averaged over subjects and parcels for each state. Power maps and coherence networks are shown relative to the average across states. The black dashed line in the state PSDs shows the time-averaged PSD. B) Inferred state probability time courses. Note, only the first 8 seconds for the first subject has been shown for illustration. C) Inferred state transition probability matrix. D) Distribution of summary statistics for dynamics across subjects.

Each network corresponds to a unique transient pattern of power and coherence. The visualisation in Figure 8A shows how the network activity, in the form of power maps and coherence networks, changes relative to the the time-averaged activity. E.g. when State 1 is active there is increased brain-wide power and parietal/temporal/frontal coherence compared to when we average over time; when State 2 is active there is increased sensorimotor power and coherence, etc. The similarity of the state power maps is quantified using the correlation in Figure S3.

The transient networks have fast dynamics with average lifetimes of less than 100 ms (Figure 8D). The probability of transitions between networks shows some structure (Figure 8C). For example, State 1 has a higher probability of transitioning to another positively activated state (States 2-4). There is also a relatively high probability of remaining in State 1 and 10 once activated, which is reflected in the higher mean lifetime for these states (Figures 8C and D).

### 3.5 Age Effects in Transient Networks

Next, characterised how the spatio-spectral and dynamic properties of the transient networks were affected by age. Figure 9 shows age effects in the power, coherence, and dynamics of each transient network.

**Figure 9:**
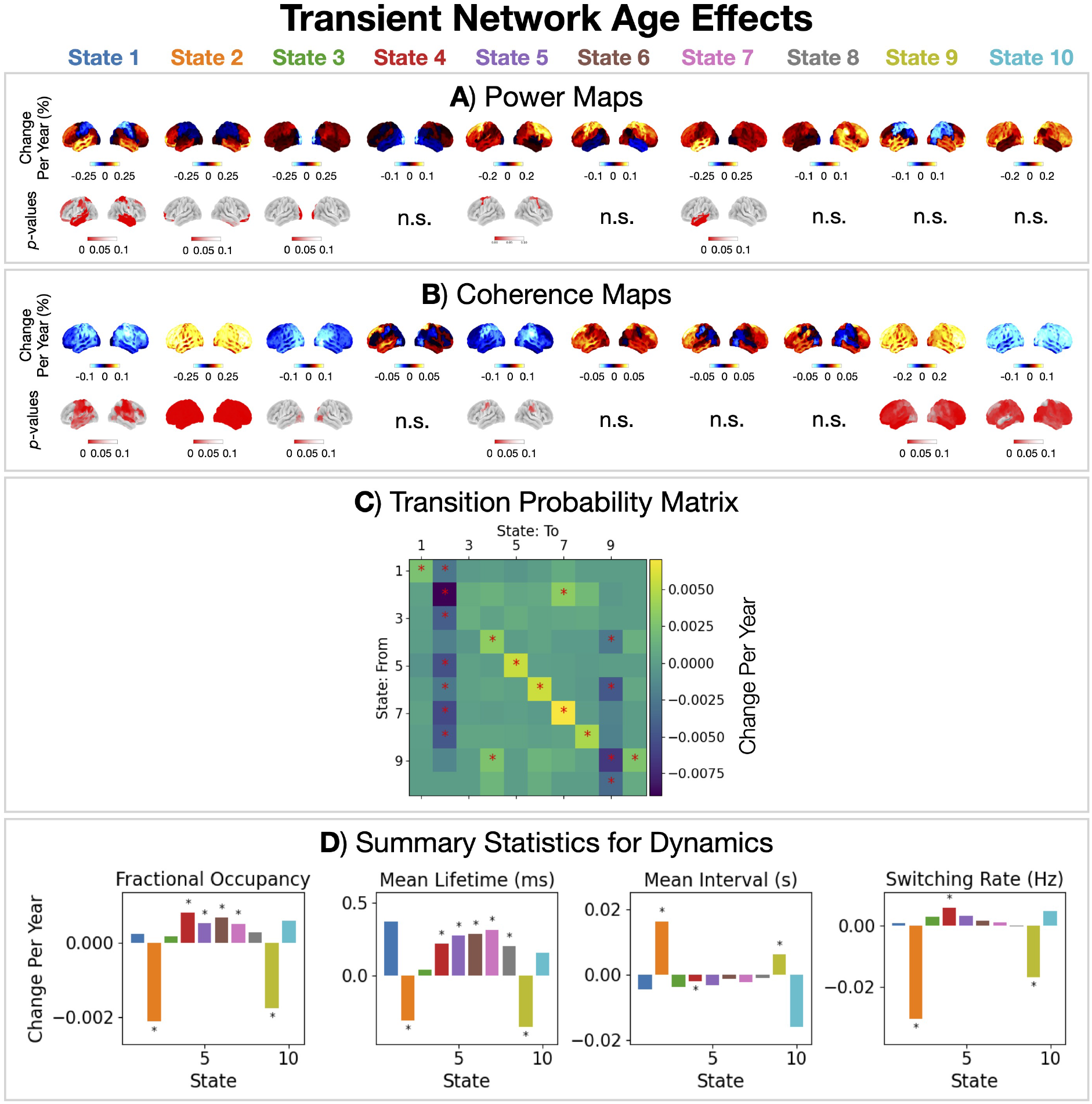
Two groups of states show opposite age effects in dynamics: frontal networks (States 2 and 9) decrease in occurrence, the others increase. Age effects in power (A), coherence (B), transition probabilities (C), and summary statistics for dynamics (D) for each HMM state. The asterisks indicate a *p*-value *<* 0.05.

There are two groups of states that show opposing age effects in dynamics. The diagonal values of the matrix in Figure 10C correspond to the effect of ageing on the *stay probability*, i.e. the probability of staying in a state, which will translate to the lifetimes each of state. Most states (1, 2, 5, 6, 7, 8 and 10) show an increase in stay probability with age, which is also reflected in the summary statistics of the dynamics (increased fractional occupancy and lifetime) shown in Figure 9D. In contrast, States 2 and 9 (frontal networks) show a decrease in stay probability, fractional occupancy and lifetime with age. The frontal networks (States 2 and 9) are particularly affected by age. Both also show an increase in brain-wide coherence (Figure 9B).

**Figure 10:**
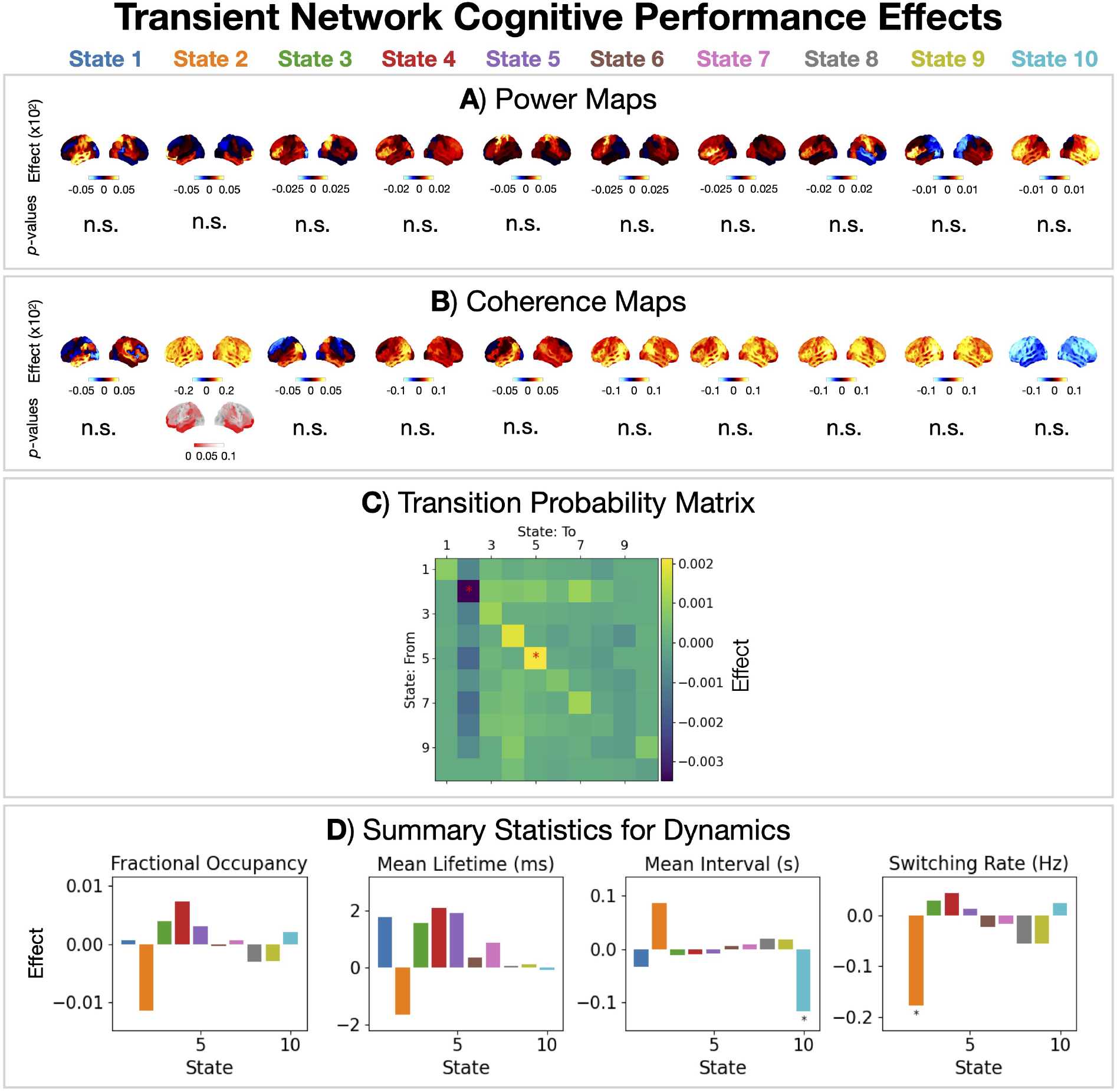
The frontal network (State 2) is linked to cognitive performance. Cognitive performance effects in power (A), coherence (B), transition probabilities (C), and summary statistics for dynamics (D) for each HMM state. The asterisks indicate a *p*-value *<* 0.05.

### 3.6 Cognitive Performance Effects in Transient Networks

The transient network perspective may provide insight into how the dynamic reorganisation of brain activity relates to cognitive function. To investigate this, we characterised the features of the transient networks that correlated with cognitive performance. Figure 10 shows the cognitive performance effects for transient networks and their dynamics.

The frontal network (State 2) shows a link with cognitive performance. We see an increase in coherence in the frontal network (Figure 10B) as well as a reduced stay probability (Figure 10C) and reduced switching into this network (Figure 10D).

## 4 Discussion

Table 2 summarises (statistically significant) age effects in time-averaged and transient functional networks. Clearly, there are many age effects in the functional brain networks of healthy individuals.

**Table 2:**
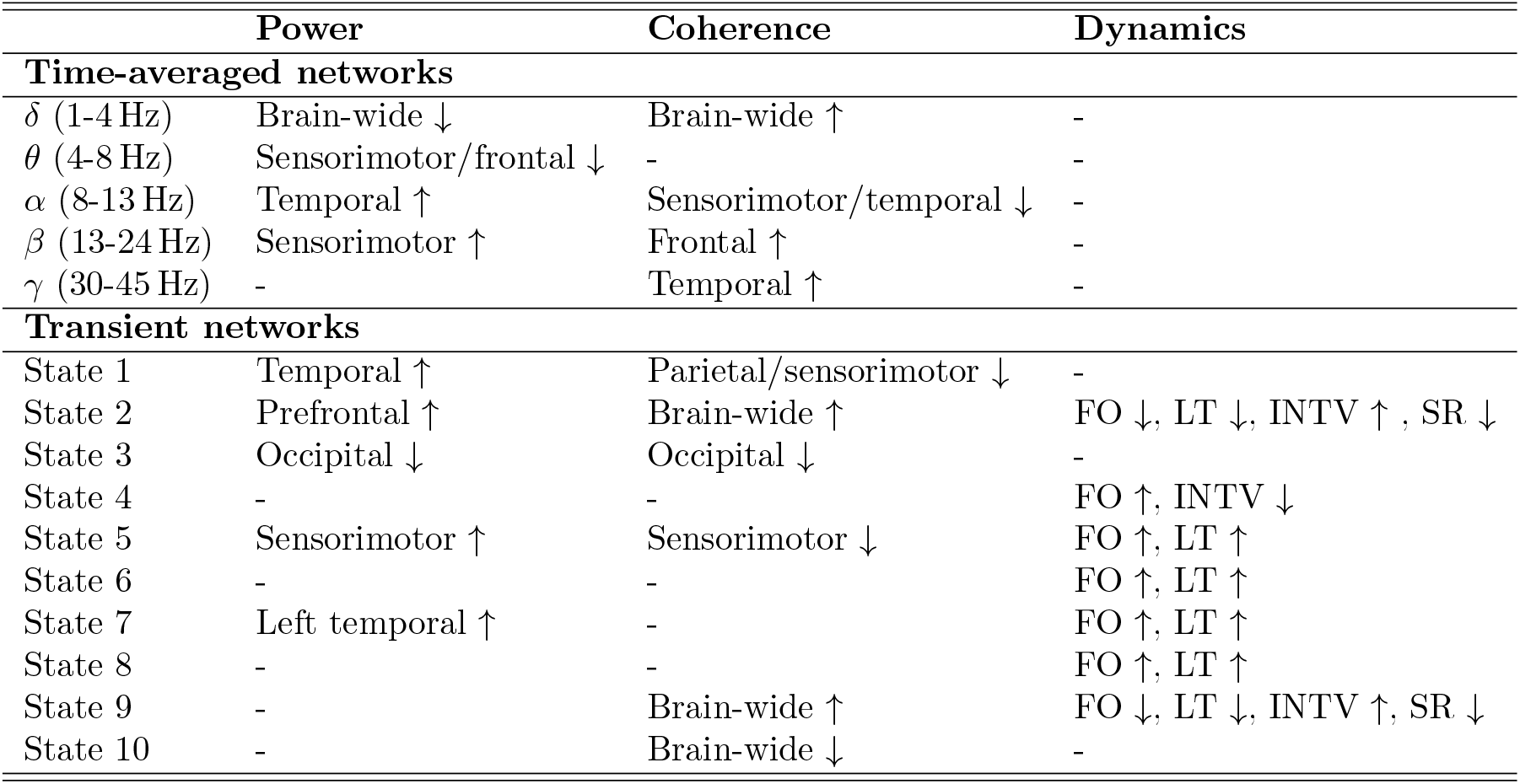
Age effects in functional networks. ↑ denotes an increase with age and ↓ denotes a decrease. Acronyms: fractional occupancy (FO); mean lifetime (LT); mean interval (INTV); switching rate (SR).

### Confounds

In this work, we included several confounds, such as brain volume, head size and position, which previously have been overlooked. These confounds and their impact on age effects are discussed in [63]. In [63], it was found that group-level age effects in MEG are robust to most of these confounds, although covariance with grey matter was found to be important to model in particular frequency bands (*δ, α, β*).

### Time-averaged networks

Multiple age effects were observed in the time-averaged power and coherence. Different changes occurred for different regions and frequencies. Relating the time-averaged networks to cognition, we found a general increase in coherence was linked to better cognitive performance.

### Transient networks

The transient network description provided a more detailed view of functional brain activity that is inaccessible in time-averaged approaches. We identified ten transient networks with unique spatio-spectral properties and fast dynamics. Multiple age effects were observed across all networks, both in the power and coherence within a network and in their dynamics. We observed age effects in the dynamics of transient networks, whereby the occurrence of the frontal networks (States 2 and 9) decreased with age, whereas all other networks increased. Relating the transient networks to cognition, we found cognitive performance was linked to the frontal network (State 2).

Below we discuss our findings in relation to previous neuroimaging and cognitive ageing studies.

### 4.1 Previous Neuroimaging Findings

#### 4.1.1 Resting-State M/EEG

##### Oscillatory power

Several studies have reported age effects in time-averaged oscillatory power. From studies that looked at linear age effects, a decrease in low-frequency power (*δ, θ*) and increase in high-frequency power (*β, γ*) have consistently been reported [47, 49, 50, 51, 52]. However, there is significant heterogeneity across studies in the regions affected. The findings in the current work support the observation that low-frequency power decreases and high-frequency power increases with age (Figure 6B). Age effects in *α*-band power are less clear with both increases and decreases previously being reported in MEG studies. This is due to the spatial heterogeneity in *α*-band power changes, similar findings were reported in [47, 48, 50]. In the current work, a decrease in occipital-*α* power due to the slowing (leftward shift) of the *α*peak frequency (Figure S1) and increase in temporal-*α* power (Figure 6B) was observed. These findings are consistently reported in the EEG [58, 59, 60, 61] and MEG [45, 46, 62, 63] literature. Note, each study covers a slightly different age range and there can be discrepancies due to sensor vs source-level analysis.

##### Time-averaged networks

Turning to age effects in time-averaged functional connectivity, Stier and colleagues [47] found a brain-wide increase in *θ* and *γ* connectivity and a brain-wide decrease in *α* connectivity with age (using imaginary coherence as the measure of connectivity). In the current work, the same findings have been observed albeit with a different measure for connectivity (coherence, Figure 6C).

##### Transient networks

Two studies have looked at transient functional networks inferred using MEG data: [56, 57]. Both used an HMM to infer the transient networks. Several of the transient networks presented in [56] are found here despite some differences in how the data were prepared for the HMM. Here, we used time-delay embedding to model dynamics in the cross spectral properties (coherence) of the data [15], whereas [56] focused on the dynamics of amplitude covariances.

Most of the age effects of the transient networks reported in [56] were also found here. However, two transient networks in [56] show opposite age effects in occurrence relative to our work. First, Tibon’s frontal network (FTP2) increases in occurrence, whereas our corresponding frontal network (State 2) decreases. Second, Tibon’s visual network (EV2) decreases in occurrence, whereas our corresponding visual network (State 8) increases. Both discrepancies arise due to differences in modelling the transient networks. In the current work, each transient network is modelled as a unique spatiospectral (i.e. amplitude and frequency) pattern of activity, whereas in Tibon’s work each transient network is modelled as a spatial (amplitude only) pattern of activity. The transient networks we identify underpin the frequency-specific age effects observed in the time-averaged power (Figure 6B). The decrease in time-averaged *δ* power with age is underpinned by a decrease in the frontal network (State 2) occurrence (Figure 9D). The decrease in time-averaged occipital-*α* power with age is underpinned by a reduction of within network power of State 8 (Figure 9A) in combination with the increased occurrence (Figure 9D). In other words, although the visual network (State 8) has a higher occurrence, the occipital-*α* power when it activates is decreased with age.

Tibon and colleagues [56] also linked the occurrence of transient networks to cognitive performance. They found better cognitive performers (in terms of fluid intelligence) had lower occurrences of frontoparietotemporal networks and higher occurrences of visual networks whilst accounting for age. Our findings support this with an increased occurrence of the visual network (State 8) and decreased occurrence of the frontal network (State 2) despite using a different metric for cognitive performance^6^ (Figure 10D).

Overall, apart from the direction of age effects in dynamics for the frontal (State 2) and visual network (State 8), this study and Tibon’s agree reasonably well in terms of the changes to dynamics with age and their link to cognition.

Coquelet and colleagues [57] have reported several results (in the Elderly − Adult group contrast) that are reproduced here. They reported an increase in time-averaged brain-wide *β* power, which we find as a sensorimotor/frontal power increase in the *β* band (Figure 6B). They used the same approach as Tibon and colleagues [56] to model dynamics in amplitude covariances to identify transient networks with an HMM. They reported an increased occurrence for the left and right auditory networks with age, which we also find (States 5 and 6, Figure 9D). They also reported a decrease in occurrence of the visual network with age (similar to the finding in [56]), which is the opposite age effect in the dynamics of our visual network (State 8). This discrepancy is for the same reason as with Tibon’s work (modelling dynamics in oscillatory properties vs amplitude). Overall, apart from the age effect of the visual network (State 8), our findings agree reasonably well with those of Coquelet and colleagues.

#### 4.1.2 Resting-State fMRI (Networks)

A well-established finding in the fMRI literature is the existence of functional networks in restingstate activity [67, 68]. The same functional networks are recruited in cognitive tasks [69, 70] suggesting that the underlying functional architecture that underpins cognition can be studied via the resting-state networks. For a review of ageing studies in fMRI resting-state networks, see [71, 72, 73, 74].

Comparing directly between the MEG and fMRI networks is difficult for a number of reasons, including: the difference in the underlying signal being measured (postsynaptic currents in MEG, BOLD for fMRI [75]); differences in the measure for functional connectivity (coherence for MEG, correlation in fMRI); and differences in the network modelling (the HMM assumes only one state is activate at a given time point, whereas the spatial ICA used in fMRI allows temporally overlapping networks [67, 68]). Nevertheless, we summarise the key findings from resting-state fMRI below and comment on the perspective provided by MEG.

##### Within and between network functional connectivity

The fMRI literature reports there is a decrease in the within-network functional connectivity with age for most resting-state fMRI networks and an increase in the functional connectivity between networks [71]. In the MEG description, we observe heterogeneous changes across the networks (Figure 9B). Whereas within-network connectivity decreases with age in States 1, 2, 3, and 10, and increases in States 2 and 9. Due to the mutual exclusivity assumption of the HMM, we do not assess inter-network connectivity in MEG.

In particular, the fMRI default mode network has consistently been reported to decrease in functional connectivity with age [71, 72, 73, 74]. In the current work, State 1 (Figure 8) most closely resembles the fMRI default mode network [13, 41] given of the overlap in regions showing high activity in MEG and fMRI (frontal and parietal). In this MEG default mode network, we also observe a brain-wide decrease in functional connectivity with age (coherence, Figure 9B).

##### Network dynamics

A reduction in resting-state fMRI network dynamics, i.e. a reduced ability to switch between network states, with age has consistently been reported [71]. In the current work, we find the rate of switching into two frontal networks (States 2 and 9, Figure 9D) are especially reduced with age, supporting the fMRI finding. Cabral and colleagues [76] related the switching rate of resting-state fMRI networks to cognitive performance (a PCA-reduced battery of cognitive test scores). Comparing older adults with good and poor cognitive performance, they found reduced switching rates and longer state lifetimes correlated with better performance. Our findings support this, despite the difference in the time scale of switching^7^. In particular, we find that a reduction in the rate of switching into the frontal network (State 2, Figure 10D) correlates with cognitive performance.

#### 4.1.3 Task fMRI

There are two main observations for age-related changes from task fMRI studies that have consistently been reported. These are discussed below.

##### Posterior-Anterior Shift in Ageing (PASA)

This is a decrease in posterior activity and increase in anterior activity observed during task for older participants compared to younger participants [66]. Although the current work has looked at resting-state MEG, there is some evidence for PASA-like age effects in connectivity (coherence). Age effects in time-averaged broadband (1-45 Hz) coherence show a clear anterior-posterior gradient (Figure 6C). However, the age effects in time-averaged power show very different spatial patterns for each frequency band (Figure 6B).

##### Task fMRI Hemispheric Asymmetry Reduction in Old Adults (HAROLD)

This is the observation that older participants show less lateralised activity in frontal regions compared to younger participants in memory tasks [64]. In the current work, we found little evidence in support of HAROLD. This may be because we are studying resting-state MEG data, although HAROLD has been reported in resting-state fMRI [65].

HAROLD suggests we should expect less lateralised activity with increasing age. A result of note in the current work is the observation that the left temporal network (State 7) shows a larger increase in fractional occupancy and mean lifetime (underpinned by the increased stay probability) with age compared to the right temporal network (State 8), see Figure 9D. The significantly increased power in the left temporal network (State 7, Figure 9A) further supports greater age effects in the left temporal lobe compared to the right. This suggests resting-state functional activity in MEG may in fact be more lateralised with age, at least in the temporal lobes, which is an observation that directly opposes HAROLD.

### 4.2 Cognitive Ageing Theories

Cognitive decline often occurs with ageing. However, there is broad variability with some individuals managing to preserve their cognitive abilities into very late life [77, 78]. Here, we studied a large cohort of individuals that show some cognitive decline across multiple domains with age (Figure 2). These individuals were deemed to be healthy by the Cam-CAN study [18, 19]. A question that arises is what aspect of brain function supports healthy cognitive ageing. Below we discuss our measure for cognitive health (the PCA-reduced cognitive score) and consider the *compensation* theory for cognitive ageing [7]. For a review of cognitive ageing theories in relation to observations from neuroimaging, see [79, 80].

#### PCA-reduced cognitive score

In this work, we reduced 10 cognitive test scores into a single measure for cognitive performance using PCA (see Section 2.1). This approach was motivated by the fact that all 10 scores were correlated, suggesting that there is a low-dimensional nature to the scores. Furthermore, it was observed that there is a single mode of variation in lifestyle, demographics and psychometric measures that corresponds to a specific pattern of brain connectivity [81]. In this work, we aimed to identify the pattern of neuronal activity that relates to a single PCA-reduced cognitive score. However, it should be noted that it is possible for the neuronal correlates of each individual cognitive test score to vary from that of the PCA-reduced score.

#### Compensation

This is the idea that individuals preserve their cognitive ability despite a failure to maintain the integrity of relevant neural resources (e.g. structural connections) by engaging some compensatory functional activity. For example, individuals might have white matter integrity that is degrading with age and a functional network that has increased activity with age. A prediction of this theory is that the cognitive performance effect should be in the same direction as age effects, i.e. those that have better cognitive performance (whilst accounting for age) appear to have biologically older looking brains in terms of their functional activity. Note, it is possible that compensatory changes can be isolated to particular functional networks rather than affect all networks equally. In fMRI studies, a common hypothesis of the frontal activity changes with age are a compensatory mechanism to counteract other age-related changes, such as structural decline, with increased functional activity [82].

We found the properties of a frontal transient network (State 2) correlated with cognitive performance and age in the same way (Figures 10 and 9 respectively), suggesting age-related changes to this network are compensatory. In other words, the changes in this functional network that are needed to improve cognitive performance are those that are happening with age. Note, we have observed this in a cross-sectional study; to conclusively demonstrate these are compensatory changes, a longitudinal study is required.

Speculatively, the age and cognitive performance effects of the frontal network could be interpreted as an increase in efficiency: less frequent, shorter visits are observed suggesting less time is needed for the frontal network to performance cognition. The reverse maybe true for the remaining networks, i.e. a reduction in efficiency indicated by more frequent, longer visits.

### 4.3 Clinical Disease Studies

In the clinical study of diseases, such as Alzheimer’s or Parkinson’s disease, age is an important confound. To understand pathological changes in the brain due to disease, we must first characterise age-related changes in the healthy brain, especially the changes expected due to healthy ageing. The current work goes towards this aim by characterising the large-scale functional networks found in a large cohort of healthy individuals. We have made these functional networks (time-averaged and transient) publicly available, see Section 6. These can be used as a point of comparison in clinical MEG studies and could potentially be a useful resource for researchers studying age-associated disease with MEG.

## 5 Conclusions

We studied the effect of age on cortical networks of oscillatory activity using MEG recordings from one of the largest cohorts to date (*N* =612, 18-88 years old) and including a comprehensive set of confounds, such as brain volume, head size and position. Our findings show many age effects should be expected for time-averaged and transient networks for healthy individuals. We showed the dynamics of the transient networks are correlated with age and are related to cognitive performance. Our results are consistent with the idea that a transient frontal network acts in a compensatory manner to preserve cognitive health with age. We have provided the networks calculated in the current work as a public resource to characterise a healthy brain, which may be useful for understanding when an individual deviates from a healthy trajectory due to disease.

## 6 Data Availability and Code

Access to the Cam-CAN dataset can be requested here [20]. Python scripts for reproducing the analysis in the current work starting from the public data are available here: https://github.com/OHBA-analysis/Gohil2025_AgeEffectsRSNs. The time-averaged and transient networks calculated in the current work are also provided.

## Footnotes

1 We use the term *effect* to refer to a regression coefficient (*β*) in a General Linear Model. This *effect* represents the association between a particular regressor (age) and the target variable whilst accounting for the influence of other confounding regressors.

2 The Cam-CAN dataset contained 643 subjects with MEG recordings, of which 22 did not have sMRIs and 9 failed in the sMRI surface extraction.

3 Also known as *spontaneous* or *task free* activity.

4 The PSD of standardised data is often referred to as the *relative PSD*.

5 Fractional occupancy, mean lifetime, mean interval and switching rate.

6 We used the PCA-reduced cognitive score, see Section 2.1.

7 fMRI network dynamics are on time scales of seconds, where MEG network dynamics are on time scales of ∼ 100 milliseconds.

