## Supplementary material for "Effects of age on resting-state cortical networks": SI

### 1 Supplementary Information (SI)

#### 1.1 Preprocessing, Source Reconstruction and Parcellation

For each subject, we applied the following preprocessing, source reconstruction and parcellation steps:

1. **MaxFilter**. The raw MEG recordings were first MaxFiltered (using v2.2.12). This is an algorithm used to remove noise originating from outside the scanner from the MEG recordings. Temporal signal source space separation (tSSS) [1] was applied (correlation threshold 0.98, 10 s sliding window) with head-motion correction (using 200 ms windows). We used the MaxFiltered MEG data (with movement compensation, without default transformation) provided by Cam-CAN.

2. **Filtering and downsampling**. First, we bandpass filtered the data (fifth-order forward-backward IIR Butterworth) between 1 and 125 Hz, then applied notch filters (2 Hz width) at 50 Hz and 100 Hz to remove mains artefacts and an additional spike artefact at 88 Hz. Following this, the data were downsampled to 250 Hz.

3. **Automated bad segment/channel detection**. Bad segments and channels can have a detrimental effect on independent component analysis (ICA) denoising (next step). Therefore, we annotated channels and segments with a significantly high variance ($p$-value $< 0.1$) as bad. For the bad channel detection we calculated the variance of each channel and determined the significance (likelihood of the channel variance being an outlier) using the generalised-extreme studentised deviate (G-ESD) algorithm [2]. Similarly, for bad segment detection, we first separated the time series in 2 s windows and calculated the variance of each window, then we calculated the likelihood of the variance being an outlier using the G-ESD algorithm[1]. The data across all channels were marked as bad if a bad segment was identified. Bad segment and channel detection was applied separately to the different sensor types (magnetometers and gradiometers). In the bad segment detection, a maximum of 10% of the time series could be marked as bad.

4. **ICA artefact removal**. After excluding the bad segments and channels, we applied FastICA [3] to the sensor-level MEG data, decomposing the signal into 64 independent components (this is the typical rank of MaxFiltered data). Based on the correlation of the source time courses with the EOG/ECG electrodes (0.9 threshold), independent components were marked as ocular/cardiac noise and removed from the MEG data.

5. **Bad channel interpolation**. Bad channels in the ICA-cleaned MEG data were replaced using a spherical spline interpolation [4]. Each sensor type (magnetometer and gradiometer) was interpolated separately. This was the final step in preprocessing the sensor-level MEG data.

[1]It is possible by removing bad segments we introduce discontinuities that artefactually give rise to state-switching dynamics. In this work, the number of bad segments identified for each subject did not correlate with HMM state switching rates.

6. **Coregistration**. Scalp, inner skull, and outer brain surface were extracted from the sMRI using FSL BET [6, 7]. The nose was not included in the scalp surface because the sMRI images were defaced. The scalp surface was then used in OSL RHINO [5] to coregister the sMRI to the MEG data based on matching the scalp to a set of digitised Polhemus head shape points and fiducials, whose positions relative to the sensors were known via the head-position coils.

7. **Forward modelling**. The *forward model* gives us the expected signal at each sensor (leadfield) for a dipole placed at a particular location inside the brain. The magnetic field from this dipole transverses different tissue types, which have different electromagnetic properties. It is common in MEG source localisation to model the changes in electromagnetic properties using a single boundary separating the volume inside the brain from outside. In the current work, we used a boundary element model [8], using the inner skull surface as the boundary. We calculated the leadfields for a dipole placed in the $x$, $y$ and $z$-direction on an 8 mm isotropic grid inside the volume bound by the inner skull.

8. **Bandpass filtering and bad segment removal**. We applied a bandpass filter to the preprocessed MEG data (firth-order forward-backward IIR Butterworth) to focus on the frequency range of interest, which is 1-80 Hz, and removed bad segments from the data to ensure these did not impact on the quality of the source localisation (next step).

9. **Volumetric beamforming**. We employed a beamformer [9, 10] to source localise the MEG data. The beamformer estimates source activity (at a location inside the brain) as a linear combination of sensor activity. In this step, we calculated the weights for the linear combination, referred to as the *beamformer weights*. We used a unit-noise-gain invariant Linearly Constrained Minimum Variance (LCMV) beamformer. This type of beamformer controls for the depth bias by normalising the beamformer weights [11].

   To calculate the beamformer weights, we need provide the leadfields from the forward model, the covariance of noise (signal measured by the sensors unrelated to brain activity) and the covariance of the data (signal measured by the sensors containing both noise and brain activity). For the noise covariance, we used a diagonal matrix containing the variance of each sensor type. We filled the diagonal elements corresponding to magnetometers (gradiometers) with the average (temporal) variance across all magnetometers (gradiometers). For the data covariance, we used the covariance of the data from step 8 reduced to a rank of 60 (this regularisation improves the robustness of the beamformer).

   The beamformer calculates the activity in the $x, y$ and $z$-direction at each grid point used in the leadfield calculation. This is reduced to a single value by projecting the $x, y, z$ activity onto the axis that maximises power of at that location. After beamforming, we are left with the activity at each grid point inside the brain, referred to as a *voxel*.

10. **Parcellation**. Often, we have a very large number of voxels, so we parcellate the data into particular regions of interest (ROIs). In the current work, we used an anatom-

ically defined 52-ROI parcellation, which is approximately symmetric. See [12] for a description of the parcellation. Parcel time courses were obtained by applying PCA to the (demeaned) voxel time courses assigned to a parcel and taking the first principal component. PCA is preferred over simply taking the mean across voxels because there is a sign ambiguity in the voxel time courses due to the beamformer.

11. **Orthogonalisation**. The source localisation of MEG data suffers from spatial leakage where the uncertainty in estimating source activity leads to nearby locations in source space exhibiting highly correlated activity. These correlations can be misinterpreted as functional connectivity (so called 'inherited connections' [13] or 'ghost interations' [14]). To substantially reduce the impact of spatial leakage, we used the symmetric orthogonalisation technique proposed in [13], which removes all zero-lag correlations from the parcel data. This can be seen as a conservative approach as we have likely removed some genuine zero-lag functional connectivity from the data.

12. **Sign flipping**. Unfortunately, there is an ambiguity in the sign of each parcel time course (due to the ambiguity of the dipole direction in the source reconstruction and due to the PCA used in the parcellation). This means the sign of each parcel time course can be misaligned across subjects. After we have the orthogonalised parcel time courses for each subject, we calculate the covariance of time-delay embedded parcel data (see Section 2.3.1) for each subject and flatten the upper triangle into a vector. Comparing the correlation of this vector for each pair of subjects, we select the median subject (highest average correlation) as a template. Finally, we match the sign of each subject's orthogonalised parcel time course to the template with the random search sign-flip algorithm described in [15] using the correlation between flattened time-delay embedded covariance matrices as the metric for alignment.

13. **Standardisation**. Finally, we temporally standardise (z-scored) each sign flipped parcel time course individually. This results in a time series with zero mean and unit variance for each parcel.

#### 1.2 Characterisation of Time-Averaged PSD with FOOOF

A popular method for characterising a PSD is FOOOF [16]. Fitting this model to the time-averaged group-level PSD in Figure 1A, we find: the aperiodic parameters are offset = -1.06 and exponent = 0.76; and we find two peaks, one in the alpha band (centre frequency = 9.2 Hz, power = 0.5, bandwidth = 4.1) and another in the beta band (centre frequency = 17.3 Hz, power = 0.3, bandwidth = 9.6).

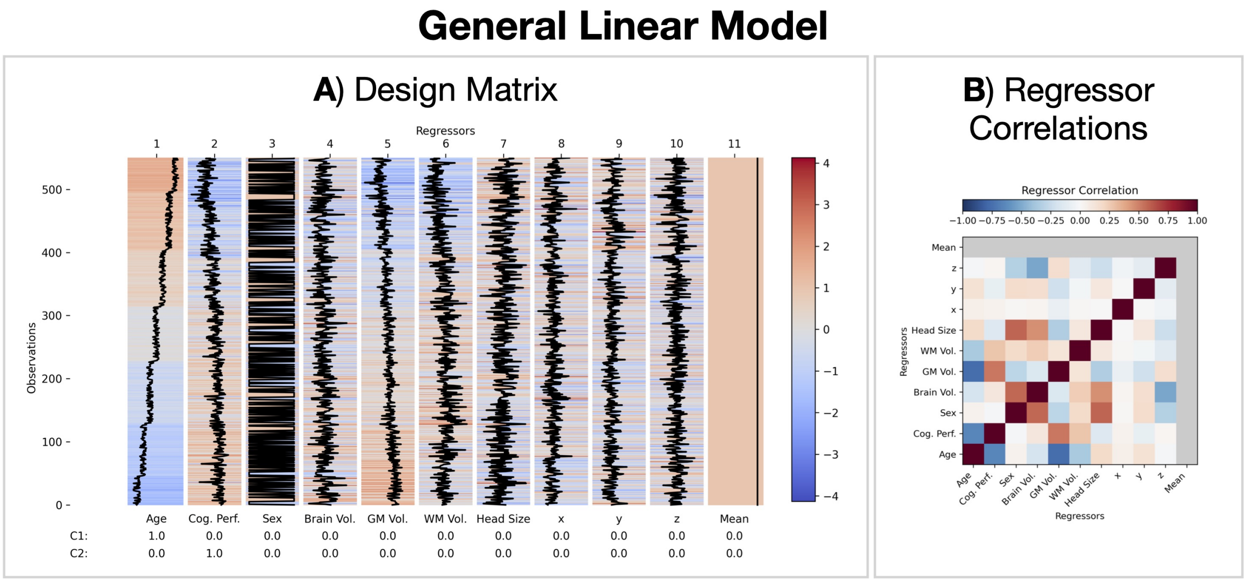


Figure S1: **General Linear Model used for group-level statistical analysis**. A) Design matrix. C1 and C2 for the Contrast of Parameter Estimates (COPEs) used to test age and cognitive performance effects for statistical significance. $x, y, z$ correspond to the central position of the head in the scanner. Acronyms: Grey Matter (GM); White Matter (WM). This design matrix was fitted to the target data stated at the start of Section 2.4. B) Correlation between regressors.

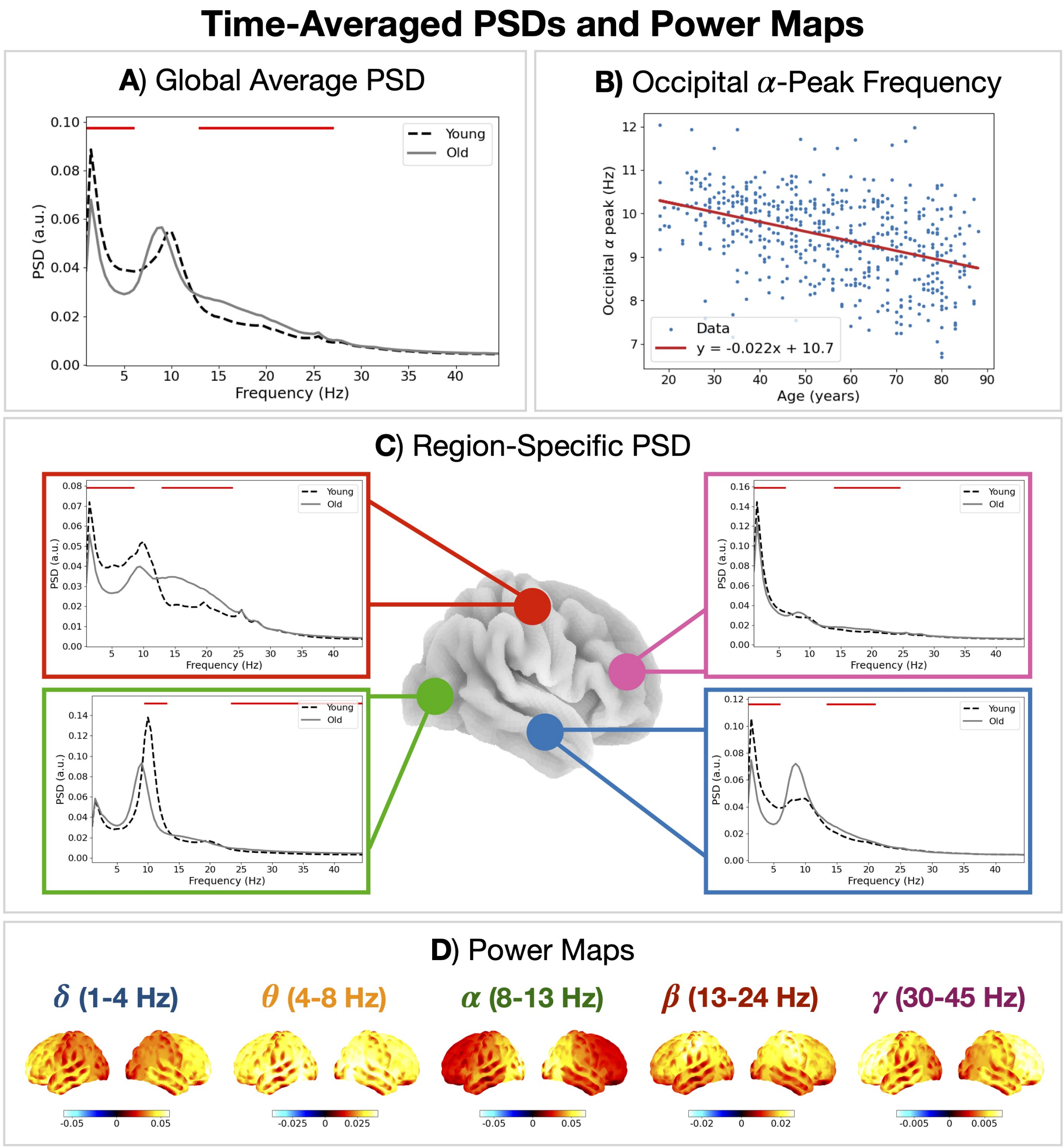


Figure S2: **Time-averaged PSDs and power maps.** A) PSD averaged over parcels and subjects for young (18-28 years old) and old (78-88 years old) participants. B) Occipital $\alpha$-peak frequency vs age. C) PSD averaged over subjects for young and old participants for different regions. The horizontal red bars indicate a $p$-value $< 0.05$ for the difference between young and old. D) Unreferenced group-averaged (all ages) power maps for each canonical frequency band.

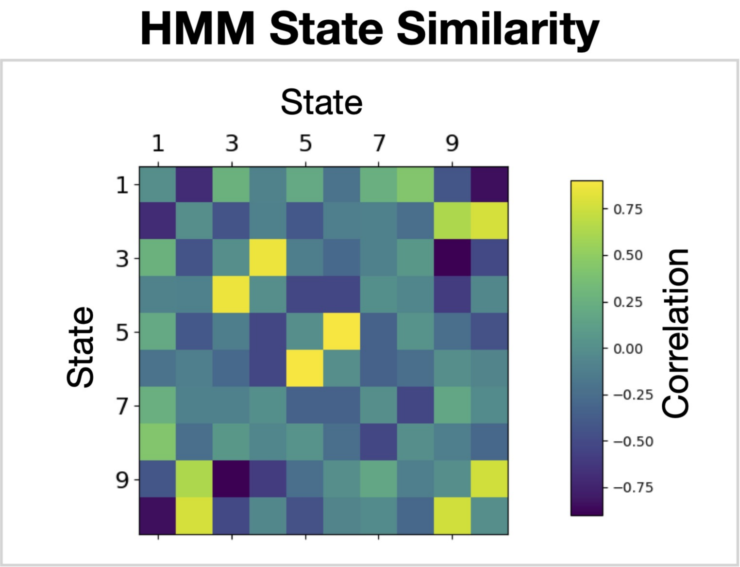


Figure S3: **HMM state similarity**. Correlation between group-level state power maps. The diagonal has been replaced with zeros.

Table S1: Hyperparameters used to train the Hidden Markov Model in `osl-dynamics`.

| **Hyperparameter** | **Value** |
|---|---|
| `batch_size` | 16 |
| `sequence_length` | 4000 |
| `learning_rate` | 0.001 |
| `optimizer` | `adam` |
| `observation_update_decay` | 0.1 |
| `trans_prob_update_delay` | 5 |
| `trans_prob_update_forget` | 0.7 |
